# Establishment and characterization of fibroblast cultures derived from a female common hippopotamus (*Hippopotamus amphibius*) skin biopsy

**DOI:** 10.1101/2020.10.14.338632

**Authors:** Tao Wang, Zelong Li, Jinpu Wei, Dongmin Zheng, Chen Wang, Chang Xu, Wu Chen, Bo Wang

**Author notes:** **Correspondence 1:** Name: Bo Wang, Address: China National GeneBank, Jinsha Road, Dapeng District, Shenzhen, China. **Correspondence 2:** Name: Wu Chen, Address: No.120, Xianlie Middle Road, Yuexiu District, Guangzhou, China.

## Abstract

The population decline in the common hippopotamus (*Hippopotamus amphibius*) has necessitated the preservation of their genetic resources for species conservation and research. Of all actions, cryopreservation of fibroblast cell cultures derived from animal biopsy is considered a simple but efficient means. Nevertheless, preserving viable cell cultures of the common hippopotamus has not been achieved to our knowledge. To this end, we detailed a method to establish fibroblast cell cultures from a female common hippopotamus fetus in this study. By combining the classic tissue explant direct culture and enzymatic digestion methods, we isolated a great number of cells with typical fibroblastic morphology and high viability. Characterization of the fibroblast cultures was carried out using different techniques. In short, neither bacteria/fungi nor mycoplasma was detectable in the cell cultures throughout the study. The population doubling time was 23.9 h according to the growth curve. Karyotyping based on Giemsa staining showed that cultured cells were diploid with 36 chromosomes in all, one pair of which was sex chromosomes. Mitochondrial cytochrome C oxidase subunit I gene sequence of the cultured cells was 99.26% identical with the *Hippopotamus amphibius* complete mitochondrial DNA sequence registered in GenBank, confirming the cells were derived from a common hippopotamus. Flow cytometry and immunofluorescence staining results revealed that the detected cells were positive for fibroblast markers, S100A4 and Vimentin. In conclusion, we isolated and characterized a new fibroblast cell culture from a common hippopotamus skin sample and the cryopreserved cells could be useful genetic materials for the future research.

## 1. Introduction

The common hippopotamus or hippo (*Hippopotamus amphibius*) is the third largest mammal on land and leads a special semi-aquatic lifestyle. They have been playing a crucial role in shaping the ecosystem they reside in. For example, these large grazers directly change the plant communities with respect to grass height and abundance and indirectly affect the abundance and diversity of aquatic animal species (McCauley et al., 2018; Stears et al., 2018). They could also influence the soil and watershed chemistry as well as silicon cycling between these two systems (McCauley et al., 2018; Schoelynck et al., 2019; Stears et al., 2018). Unfortunately, this megaherbivore population has undergone a dramatic decrease in number as a result of illegal hunting and habitat degradation (Lewison, R. & Pluháček, 2017). Long-term storage of different genetic resources is a powerful method to facilitate animal species conservation. Indeed, Saragusty et al. made an effort to cryopreserve epididymal sperm of the common hippopotamus (Saragusty et al., 2010). To our knowledge, however, preserving other types of genetic resources from the common hippopotamus is rarely reported.

Fibroblast cells are a type of mesenchymal cells that synthesize collagen and remodel extracellular matrix framework to maintain the tissue structural integrity *in vivo*. They are also valuable genetic materials for *in vitro* experiments. For instance, fibroblast cells could be reprogrammed into pluripotent stem cells for potential applications (e.g., Mishra et al., 2014; Takahashi, 2006; Verma et al., 2012). Besides, these cells were widely applied as donor cells in animal cloning and somatic cell nuclear transfer technique (e.g., Liu et al., 2018; Ruan et al., 2019; Zhai et al., 2018). Till now, generation of fibroblast cell cultures from animal species particularly those endangered ones has been frequently accomplished. Noteworthily, a majority of reported fibroblast cell cultures were derived from skin samples (e.g., Mishra et al., 2014; Siengdee et al., 2018; Wang et al., 2020). The reason lies in the fact that fibroblast cells take up a large proportion of the cellular components within the dermis and sampling a small piece of skin tissue is unlikely to cause severe damage to the animals. The skin biopsy is thus regarded as an ideal sampling site for culturing fibroblast cells *in vitro*.

Tissue explant direct culture and enzymatic digestion were two methods that were generally taken advantage of for primary cell culture (Mestre-Citrinovitz et al., 2016). For the former method, collected samples were excised into small explants which were later seeded into culture flasks to enable outgrowth of the cells. For the latter one, tissue disassociation by incubation with trypsin or collagenase was involved and acquired cell suspension was then transferred to the culture flasks. And yet over-trypsinization could probably have an adverse impact on the subculture, while comparatively collagenase digestion is considered a preferable choice (Nanda et al., 2014; Su et al., 2015). Explant direct culture without enzyme treatment could better keep the integrity of the cells and provide some essential growth factors for cell culture as well (Hendijani, 2017). But the disadvantage of this method is that it usually takes much longer to complete the primary culture especially when several rounds of cell migration from the explants were carried out to harvest as many cells as possible. In comparison, enzymatic digestion method could accelerate the process of primary cell culture significantly. It might be achievable to obtain more cells within a short culture period by combining these two abovementioned methods together.

In this study, we detailed a method of isolating a large quantity of fibroblast cells with high viability and featured morphology from a common hippopotamus fetus skin. We also adopted complementary techniques to characterize the cultured cells. We believe this newly-established cell culture could become a useful biological resource to help preserve common hippopotamus genetic diversity and facilitate future research.

## 2. Materials and methods

### 2.1 Sample collection

An adult common hippopotamus gave birth to a female calf in Guangzhou Zoo but unfortunately the fetus died very soon thereafter. The skin tissue of the dead fetus was then collected and delivered to the laboratory in Dulbecco’s Modified Eagle’s medium (DMEM) medium (Gibco, 10569-010) supplemented with Penicillin/Streptomycin (P/S) (500 U/mL) (Gibco, 15140122) and 1% amphotericin B (Gibco, R01510) at 4°C. All procedures involving animals in our study were in accordance with the ethical principles and guidelines of the institution and approved by the Institutional Review Board on Bioethics and Biosafety of China National GeneBank (CNGB) (NO. CNGB-IRB 2001).

### 2.2 Primary culture and subculture

Primary cell culture was performed using tissue explant culture and enzymatic digestion methods with a few modifications. Briefly, a thin layer of adipose tissue underneath the skin was removed carefully in the first place. The skin biopsy was then immersed in sterile saline solution supplemented with P/S (10,000 U/mL) for 10 min and minced into small explants measuring 1 cm^2^ in size. After washed three times with saline solution supplemented with P/S (1000 U/mL) and 2% amphotericin B, the explants were cut into smaller ones, say, with 1 mm^2^ in size. The tissue explants were then collected into falcon tubes and washed gently with saline solution for three times. Finally, the small explants were seeded into 75-cm^2^ cell culture flasks (Thermo Scientific, 156499) and added with pre-warmed complete culture medium I consisting of DMEM/F12 (Gibco, C11330500BT), 20% fetal bovine serum (FBS) (ExCell, FSP500), P/S (200 U/mL) and 1% amphotericin B for primary culture. The complete culture medium I was exchanged every 3-4 d before subculture.

The tissue explants were re-collected into a sterile falcon tube when the cell confluence reached ~90%. Meanwhile, the cell culture was added with 0.25% trypsin solution (Gibco, 25200072) and incubated at 37°C shortly. When the cells were observed round under a light microscope (Olympus, IX73), pre-warmed complete culture medium II consisting of DMEM/F12, 20% FBS, P/S (100 U/mL) and 1% amphotericin B was supplemented to deactivate trypsin. Cell suspension were obtained by pipetting the culture medium softly, followed which centrifugation and resuspension in new culture medium II were performed. The collected cells were then divided into new cell cultures with the initial density of 1.0 × 10^4^ cells/cm^2^. On the other hand, the re-collected tissue explants were incubated with collagenase IV solution (Gibco, 17104019) at 37°C in a shaker. After ~1 h of the digestion process, the remained tissue explants were pipetted repeatedly for acquirement of the cell suspension. After centrifugation and resuspension in fresh culture medium II, the cell suspension was transferred to culture flasks for subculture. All cell cultures were kept in a humidified incubator with 5% CO_2_ at 37°C.

### 2.3 Cryopreservation and recovery

Cultured cells were collected upon reaching cell confluence and cell viability was measured based on trypan blue staining method. The cells were then resuspended in freezing medium composed of 90% complete culture medium II and 10% DMSO (Sigma, D2650-100ML) to reach a final concentration of 2.0 × 10^6^ viable cells/mL. Division of cell suspension into cryogenic vials was performed with each vial containing 1 mL in total. The cryogenic vials were kept in a Nalgene Cryo 1°C Freezing Container (Thermo Scientific, 5100-0001) in a −80°C refrigerator overnight, after which they were stored in liquid nitrogen for long.

Recovery of the frozen cells was conducted in the water bath at 37C quickly and warmup was completed when little ice in the cryogenic vials could be observed. The thawing cells were re-evaluated regarding its viability and then resuspended in fresh complete culture medium II for subsequent culture as mentioned above.

### 2.4 Microorganism detection

Primary detection of microorganism contamination was carried out via continuous observations of the cell cultures under a light microscope. To further test whether there was bacteria or fungi growth in the cultures, BacT/ALERT 3D Microbial Identification System (BioMérieux) was employed. In order to detect the presence of mycoplasma in cell cultures, nested PCR was performed using PCR Mycoplasma Detection Set (Takara, 6601).

### 2.5 Growth curve analysis

To study the cell growth properties and calculate the population doubling time (PDT), we reseeded the cells of passage 3 (P3) into a 96-well culture plate with a concentration of 5.0 × 10^4^ viable cells/mL. A total of 5.0 × 10^3^ cells were inoculated per well to initiate the culture and six replicates for everyday measurement were used. To evaluate the cell growth, the cultured cells were supplemented with WST-8 (Beyotime, c0038) and incubated for another 1 h before the following detection. The culture plate was then put into a microplate reader (Thermo Scientific, Multiskan GO) for inspection of the absorbance value at 450 nm and this optical density (OD) value could indicate the relative concentration of viable cells. A growth curve was thereafter plotted with culture time on the X axis and Napierian logarithm of OD value on the Y axis using GraphPad Prism 7.0. PDT was calculated according to this formula: PDT = T×ln2/ln(OD_7_/OD_0_), where T is the total culture period, OD_7_ is the cell density on the 7th day and OD_0_ is the initial cell density.

### 2.6 Karyotyping

Karyotyping was performed based on a previously-reported protocol with a few modifications (Bates, 2011). Briefly, reseeded cells of P4 were cultured for 48 h after which colcemid solution (JISSKANG, HYT9087) was added with a final concentration of 1 μg/mL. After 6 h of treatment, the cells were harvested and incubated with pre-warmed hypotonic solution—0.075 M KCl—at 37°C for 10 min. Cold fixative solution consisting of methanol and glacial acetic acid in a ratio of 3:1was added to the cell pellets after centrifugation. The fixation procedure was repeated three times. The cells in fixative solution were then dropped onto clean cold glass slides from a height of ~15 cm. Afterwards, the slides were aged for 2 h in a dryer at 85°C and stained with Giemsa solution (SinoChrome, CM-G-250) according to the instructions. Imaging under a light microscope was carried out and ~20 cell spreads were analyzed using Photoshop CC 2019.

### 2.7 Mitochondrial cytochrome C oxidase subunit I (*COI*) gene sequencing

Genome DNA from the collected cells of P2 was extracted under the instructions of the user’s manual for TIANamp Genomic DNA Kit (TIANGEN, DP304-02). PCR was later conducted to amplify mitochondrial *COI* gene fragment using the primer pair as follows: VF1d_t1: 5’-TGTAAAACGACGGCCAGTTCTCAACCAACCACAARG AYATYGG-3’; VR1d_t1: 5’-CAGGAAACAGCTATGACTAGACTTCTGGGTGG CCRAARAAYCA −3’. The products were sequenced on an ABI3730 DNA sequencer and the sequence data were deposited in the CNGB Sequence Archive (CNSA) (https://db.cngb.org/cnsa/) (Guo et al., 2020) of China National GeneBank DataBase (CNGBdb) (Chen et al., 2020). Nucleotide BLAST program was run to compare the sequence identity of mitochondrial *COI* gene we submitted with those already deposited in GenBank.

### 2.8 Flow cytometry analysis

A total of 2.0 × 10^6^ cells of P3 were incubated with fixation buffer (eBioscience, 88-8824-00) for 20 min and subsequently with permeabilization buffer (eBioscience, 88-8824-00) for another 20 min. After incubated with blocking buffer—10% goat serum albumin in PBS—for 15 min, the cells were resuspended in PBS for subsequent labeling. Collected cells were then incubated with the antibodies against Alexa Fluor® 488-conjugated anti-S100A4 (1:250, Abcam, ab196380) or FITC-conjugated anti-Vimentin (VIM) (1:25, Abcam, ab128507) for 30 min. Meanwhile, incubation of the cells with Alexa Fluor® 488-conjugated IgG antibodies (Abcam, ab199091) was conducted as isotype controls. All incubation procedures were performed at room temperature in the dark and washing procedures with clean PBS after each incubation step was repeated three times. Following the last incubation and washing step, cell suspension was transferred into fluorescence activated cell sorting tubes. A minimum of 1.0 ×10^4^ cells for each sample was loaded on a BD FACSJazz cell sorter and FlowJo 10 was applied for data collection and analysis.

### 2.9 Immunofluorescence staining

Cultured cells of P2 in a 24-well plate were incubated with fixation buffer for 15 min at room temperature, followed by incubation with permeabilization buffer at room temperature for 1 h. After blocking with 10% goat serum albumin in PBS for 1 h, incubation with antibodies against Alexa Fluor® 488-conjugated anti-S100A4 (1:250) or FITC-conjugated anti-VIM (1:25) was performed in the dark at 4°C overnight. Afterwards, incubation of the cells with DAPI (Beyotime, C1005) in the dark was carried out at room temperature for 20 min. The washing procedure between incubation steps were the same as described above. Imaging of the immunostained cells was performed under a fluorescence microscope (Leica, DM4000B).

## 3. Results

### 3.1 Isolation and cryopreservation of the skin fibroblast cells

In this study, we combined the classical tissue explant direct culture method and enzymatic digestion method to obtain a great number of skin biopsy-derived fibroblast cells. In detail, elongated and spindle-like cells were observed to emigrate from the skin tissue explants rapidly (Figure 1A). Besides, the cells were tightly adherent to the plastic but sensitive to the trypsin treatment. Those properties indicated that the cultured cells were of fibroblastic origin. It was noteworthy that the cells acquired from the collagenase digestion of the re-collected tissue explants exhibited quite similar features with those migrating directly from the explants. In this way, we could isolate a large number of cells from the skin sample within a relatively short period. This new fibroblast cell culture was named CNGBCCAC00088.

**Figure 1.**
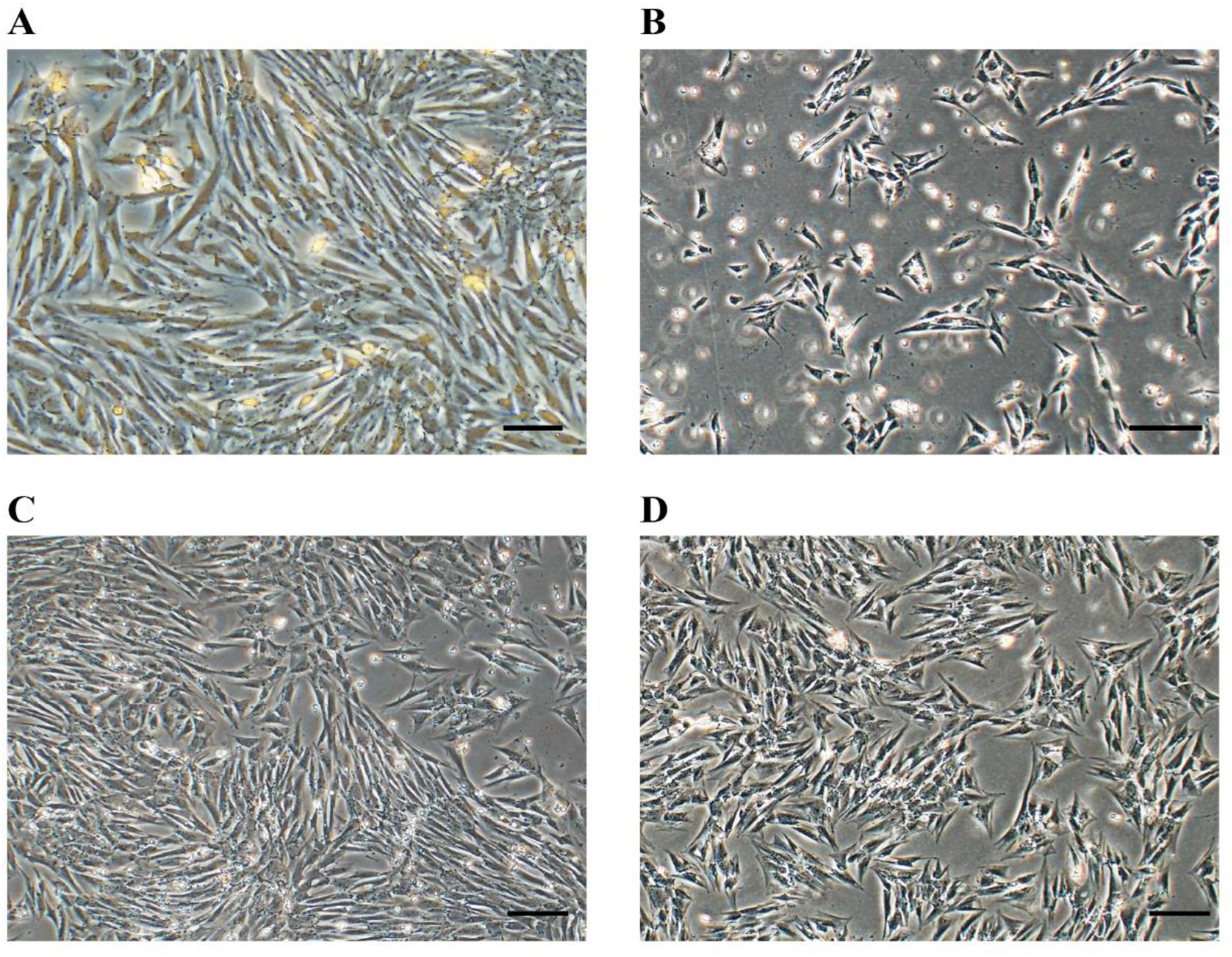
Morphology of the isolated fibroblast cells. **A.** After 6 d of primary culture, substantial cell outgrowth from the skin tissue explants collected from a newborn female common hippopotamus could be observed. The cells were tightly adherent to the plastic and spindled-shaped. Scale bar = 100 μm. **B.** 3 d after thawing and reseeding, adherent cells of P2 exhibited a long-thin and multipolar appearance indicating they were in a healthy state. Scale bar = 200 μm. **C.** Reseeded cells of P2 proliferated rapidly and aligned in a vortex-like shape after 7 d of culture. Scale bar = 200 μm. **D.** Fibroblast cells of P4 were of a normal morphology and of fast growth. No dramatic sign of cellular senescence, such as enlarged cellular body and vacuoles, was observed. Scale bar = 200 μm.

Passaged cells showed a characteristic morphology of fibroblasts and proliferated fast (Figure 1B). 7 d after replating, cell confluence reached ~70% (Figure 1C). It was apparent that cells were crowded together to display a vortex-like arrangement, which demonstrated again their fibroblastic origin. When subcultured continuously *in vitro*, fibroblast cells of P4 did not show dramatic changes in their cellular size, shape as well as proliferation rate (Figure 1D). Harvested cells of P1 and P2 before cryopreservation had viability with 90% and 93%, respectively. Noteworthily, the cell viability for different passages remained higher than 90% after recovery. These results indicated that the cell cultures were in a healthy state.

### 3.2 Detection of microorganism contamination

The presence of microorganisms could lead to a failure of cell culture and therefore we detected whether microbial contamination was extant in our established cell cultures. Not surprisingly, no actively moving bacteria or branched mycelium of fungi was found through direct observations under a light microscope. And there was no rapid onset of culture medium turbidity, which further indicated no bacterial proliferation occurred. Negative results obtained from the microbial identification system were consistent with the abovementioned findings. Additionally, no amplified DNA products from nested PCR were observed excluding the possibility of mycoplasma contamination.

### 3.3 Growth curve analysis

So as to find out the growth kinetics of the isolated cells, we recorded everyday cell density for a whole week. The plotted growth curve showed that the cell concentration increased exponentially at the beginning and reached its maximum on the 3^rd^ day (Figure 2). Cell number started to decline on the 4^th^ day and stayed relatively steady onwards till the culture ended. Based on the formula as described in the methods, PDT was 23.9 h for this common hippopotamus fibroblast cell culture.

**Figure 2.**
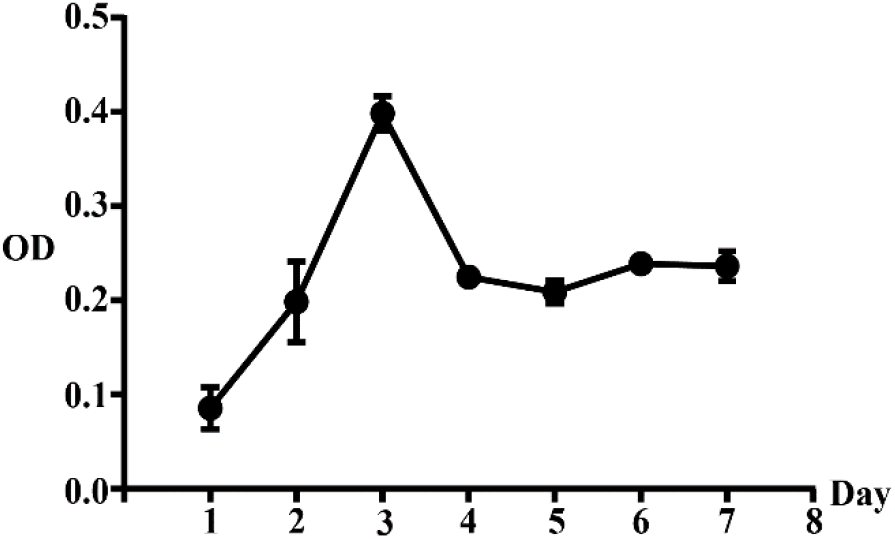
Growth curve of the fibroblast cell culture. Fibroblast cells of P2 were plated and measured regarding the cell density on a daily base. A growth curve was plotted based on the 7-d continuous measurements of OD values. PDT was calculated to be 23.9 h according to the mathematical formula described in the method. Data are plotted as Mean ± SD.

### 3.4 Chromosome analysis

It is reported that the common hippopotamus is diploid with 36 chromosomes in total (Arrighi, 2006). We collected cultured cells of P4 and performed karyotyping to verify previous results. Figure 3A is a representative of the scattered chromosomes derived from a single fibroblast cell. It could be observed that all of the chromosomes were metacentrics. An example of the chromosome pairing was presented in Figure 3B. In our study, almost every analyzed cell contained 18 pairs of chromosomes in all except one missing a chromosome. That is to say that the cultured cells contained 17 pairs of autosomes and one pair of X chromosomes, which is in conformity with the previous study. The result further clarified that the isolated fibroblast cells were indeed originated from a female common hippopotamus rather than a male one.

**Figure 3.**
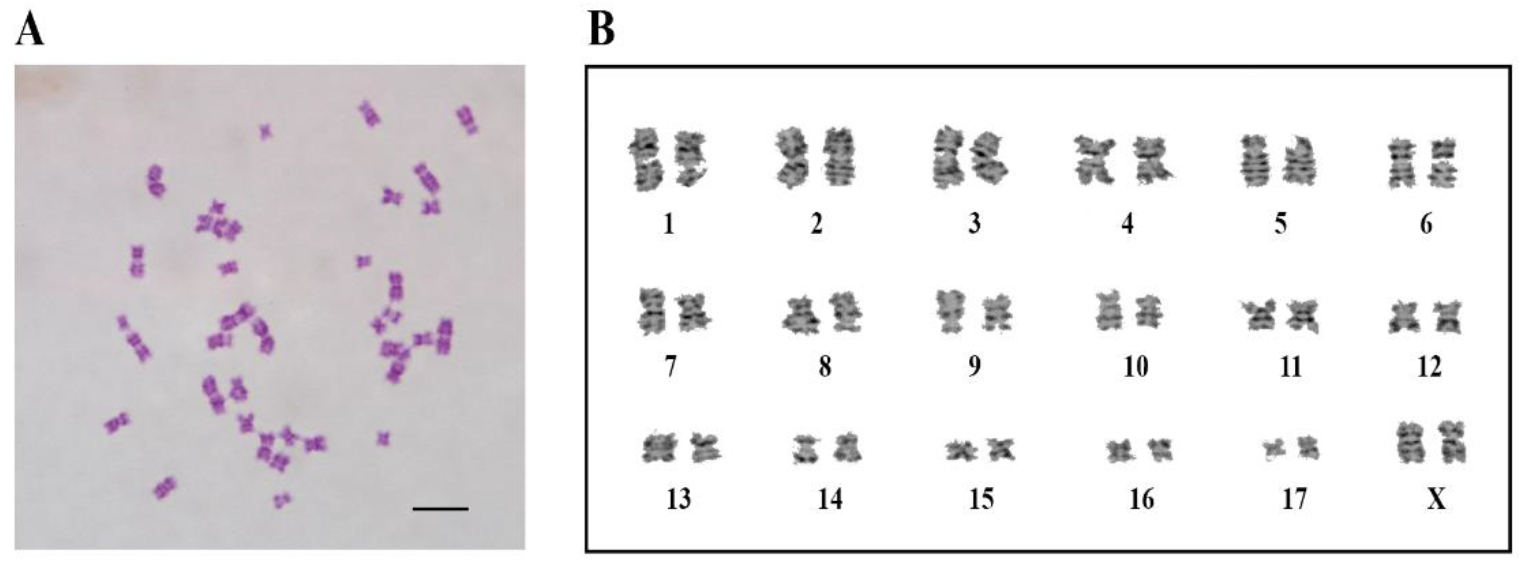
Karyotyping of the established fibroblast cell culture. **A.** Karyotyping of the fibroblast cell culture of P4 was performed based on Giemsa staining method. All chromosomes were metacentrics. A representative of the cell spreads is shown here. Scale bar = 10 μm. **B.** On the grounds of the G-banding, a total of 36 chromosomes were rearranged into 18 pairs, one of which was X chromosomes.

### 3.5 Analysis of mitochondrial *COI* sequence

In order to confirm the derivation of the isolated cells from the common hippopotamus, the mitochondrial *COI* gene was amplified and sequenced. The sequence data and associated detailed information are now available in CNSA of CNGBdb with project code CNP0001211 (https://db.cngb.org/search/sequence/N_000000918.1/). In detail, the acquired *COI* fragment was 681 bp in length with GC content accounting for 46.4%. Alignment result based on Nucleotide BLAST program running illustrated that the amplified *COI* region in our study were of 99.26% identity with the *Hippopotamus amphibius* complete mitochondrial DNA sequence registered in GenBank (Accession number: AJ010957.1). Our *COI* gene sequence was also 98.97% identical with another submitted sequence from the species of common hippopotamus (Accession number: AP003425.1). It could be concluded that the cultured cells were authentically derived from a common hippopotamus.

### 3.6 Analysis of protein expression patterns

We adopted flow cytometry and immunofluorescence staining method to identify protein expression patterns of the isolated cells. S100A4 and VIM are two well-recognized markers for fibroblasts and thus they were applied here to characterize the harvested cells. Flow cytometry analysis showed that the cells were strongly positive for S100A4 but weakly positive for VIM (Figures 4A-B). Detailed, 91.2% of the detected cells were positive for S100A4, while only 11.0% of the cells were positive for VIM. The protein expression patterns were further clarified using the immunofluorescence staining method (Figure 4C). It is worth noting that results from these two methods were a little inconsistent with respect to the VIM expression. In other words, the latter method showed that a greater percentage of the cultured cells were positive for VIM (Figure 4C). The short incubation time (30 min) with antibodies against anti-VIM in the flow cytometry method might account for this discrepancy. Nevertheless, the established cell cultures positive for fibroblast markers could demonstrate again their fibroblastic properties.

**Figure 4.**
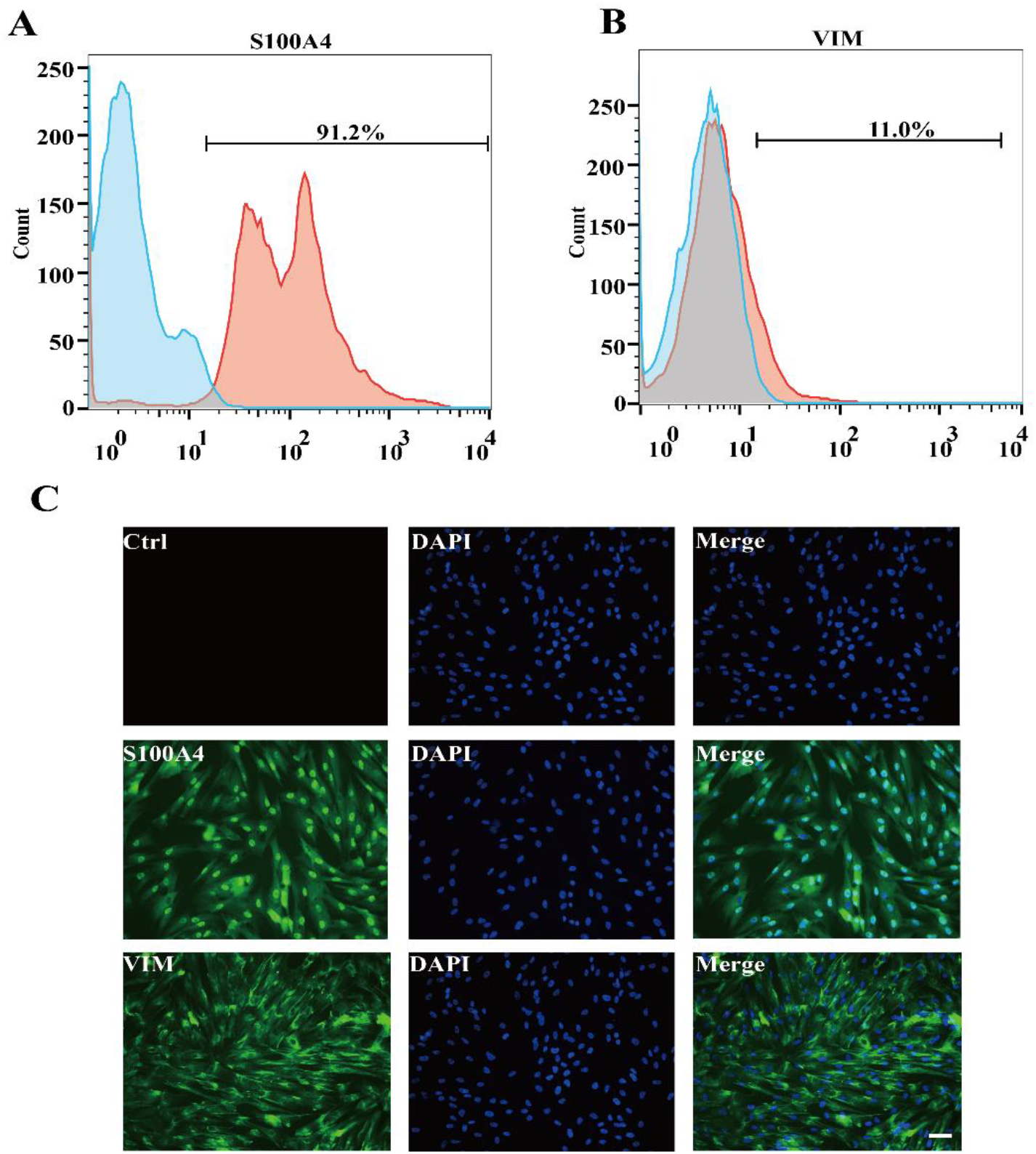
Protein expression pattern analysis of the cultured cells. **A-B.** Cells of P3 were incubated with antibodies against Alexa Fluor® 488-conjugated anti-S100A4 or FITC-conjugated anti-VIM and then analyzed by flow cytometry. Results showed that collected cells were strongly positive for S100A4 but weakly positive for VIM. **C.** Immunofluorescence staining of the cultured cells of P2 with anti-S100A4 antibodies and anti-VIM antibodies was performed separately. It could be observed that the cells exhibited high-level expression of both fibroblast markers— S100A4 and VIM. Scale bar = 100 μm.

## 4. Discussion

The common hippopotamus is native sub-Saharan megaherbivores and important part in the local ecosystem. Due to illegal hunting and substantial habitat loss, this large mammal population has been facing a rapid decline. Among multiple activities of maintaining the common hippopotamus, advanced cryopreservation technique is an appealing means to conserve the genetic diversity (Siengdee et al., 2018). In this study, we described a protocol in detail to establish high-quality fibroblast cell cultures from a female common hippopotamus fetus skin sample.

Generally, washing the collected samples with saline solution containing antibiotics was conducted at the very beginning of the primary culture (Siengdee et al., 2018; Wang et al., 2020). Previous studies confirmed the effectiveness of adding more antibiotics and washing steps on alleviating microbial contamination in the cell culture (Yap et al., 2019). Considering the sample was collected in an inhospitable environment and there was a high chance for the contamination, we increased the antibiotics concentration in the saline solution and added additional washing steps. In addition, the initial complete culture medium was supplemented with more antibiotics as well. It turns out that our modified protocol efficiently prevented the microbial growth in the cell cultures and no detectable contamination was found throughout the whole study. Meanwhile, higher concentration of antibiotics did not significantly cause adverse effects on cell properties.

To initiate the primary culture, tissue explant direct culture and enzymatic digestion methods were combined in our study. We firstly employed the former method to obtain the first batch of cell cultures and then the latter one to digest the re-collected explants. Trypsin has been reported to have a very harsh effect on the cell membrane proteins and is more likely to cause cell lysis in the suspension (Reichard and Asosingh, 2019; Su et al., 2015). Therefore, collagenase was our option to dissociate the skin explants and produce single cell suspension. In this way, we obtained a large quantity of fibroblast cells with high viability. Alternatively, Shahini et al. reported another approach to yielding enough cells from the muscle sample (Shahini et al., 2018). In that study, they started a couple of more rounds of tissue explant seeding in the culture dishes upon cells reaching confluence. Although that protocol could take full advantage of the tissue explants, the whole cell outgrowth process took a longer period. Anyhow, we think our method could enable us to acquire a large number of high-viability cells and shorten the process of primary culture.

We employed several techniques to characterize the isolated cells. The observations on the morphology and protein expression patterns indicated that the cultured cells were fibroblastic. The karyotyping and mitochondrial *COI* sequence analysis verified the fibroblast cell cultures were authentically derived from the common hippopotamus instead of other animal species. It should be noted that all experiments were conducted using cells of early passages. Not like immortalized cells, fibroblast cells have a limited number of replications and become senescence after continuous passaging *in vitro* (de Magalhães and Passos, 2018). We do not exactly know how many times our established cell cultures could be subcultured before they display dramatic signs of cellular senescence. It might be helpful to figure out when the senescence occurs by passaging the cells serially *in vitro* in the future study. Further characterization of cultured cells at different passages might be necessary to determine their phenotypic stability.

## 5. Conclusions

In this study, we detailed a method to establish and characterize fibroblast cell cultures from the skin biopsy of a newborn female common hippopotamus. In brief, additional washing of the collected samples and higher concentration of antibiotics in the medium were applied to prevent microorganism contamination. Tissue explant direct culture and collagenase digestion were combined to enable obtainment of a larger number of viable cells within a relatively short period. As a matter of fact, the isolated cells displayed typical morphology and growth property of fibroblasts. Besides, the cells were of high viability and free of microbial contamination. Characterization of the cell cultures further confirmed their fibroblastic features and origination of animal species. We believe this high-quality cell culture could be a useful biological resource for conserving genetic diversity of the common hippopotamus and facilitating future scientific studies.

## List of abbreviations

CNGB: China National GeneBank
CNGBdb: China National GeneBank DataBase
CNSA: CNGB Sequence Archive
COI: Cytochrome C Oxidase Subunit I
DMEM: Dulbecco’s Modified Eagle’s medium
FBS: Fetal Bovine Serum
OD: Optical Density
PDT: Population Doubling Time
P/S: Penicillin/Streptomycin
P3: Passage 3
VIM: Vimentin

## 6. Acknowledgement & Fund

This research was supported by the fund from the Shenzhen Key Laboratory of Environmental Microbial Genomics and Application (No. CXB201108250097A).

